# Insights into restlessness as a potential novel indicator of impaired welfare in bulls fattened for meat production

**DOI:** 10.64898/2026.03.29.715061

**Authors:** Sara Hintze, Theresa Wildemann, Florian Krottenthaler, Christoph Winckler

## Abstract

Restlessness is a symptom of chronic boredom in humans and a behavioural phenomenon anecdotally described as a concern in bulls raised for fattening purposes, but it has so far not been addressed in research. The two studies presented in this paper aimed to gain first insights into restlessness in bulls. We operationally defined restlessness by the number of transitions between behaviours in a given time period, and quantified restlessness in bulls of different weight classes (300, 400, 500 kg) on farms keeping bulls on fully-slatted floors (n=8, Study 1) as well as across three different husbandry systems (fully-slatted floor (FS, n=4), straw-based (SB, n=4) and organic pasture (OP, n=3), Study 2). All farms were visited twice, and the behaviour of different individuals was continuously recorded for 15 minutes each between 9 a.m. and 5 p.m. (Study 1) and for 8 minutes each between 6 a.m. and 10 p.m. (Study 2). The effects of weight class and husbandry system were analysed using generalised linear mixed-effects models, and we ran a sequence analysis to cluster observations by the sequence, frequency, and duration of bulls’ behaviours in Study 1. Bulls kept in fully-slatted floor systems in Study 1 changed their behaviour on average 48.3 times per 10 minutes, with high variability both within and across farms. Weight class did not have a statistically supported effect on the number of transitions, and the sequence analysis revealed four clusters that differed in sequence and in the number of transitions. In Study 2, OP bulls showed fewer transitions than SB and FS bulls (X^2^ = 23.6, p < 0.001), while SB and FS bulls did not differ. While SB pens were more structured and offered more space per animal, both SB and FS systems can be characterised by monotony, which may explain the similar level of restlessness in both systems. Alternatively, or in addition, the high feeding intensity in SB and FS systems may have led to more transitions than in the OP system, potentially due to subacute ruminal acidosis and/or laminitis and the resulting pain. However, given the study’s descriptive nature, these explanations are speculative and require systematic disentanglement in future studies. Providing initial insights into restlessness in bulls, this study lays the groundwork for future research to address the causes underlying restlessness and investigate its association with bull welfare.

**Highlights:**

- Operationally defining restlessness: number of behaviour transitions per time unit
- We quantified restlessness in fattening bulls across different husbandry systems
- Less restlessness observed in bulls kept on pasture than in indoor systems
- No difference in restlessness in fully-slatted floor and straw-based systems
- Restlessness may result from chronic boredom, intense diet or their combination

## 1 Introduction

Bulls fattened for meat production are often housed in intensive systems. Such systems are typically characterised by barrenness, monotony, high stocking densities, and fully-slatted floors, affecting both physical health, for example, carpal joint swelling on concrete slats (e.g., Brscic et al., 2015a), and behaviour, for example, resting (e.g., Gygax et al., 2007). In addition to the housing conditions, high-concentrate diets are a risk to bull welfare as they increase the risk of subacute ruminal acidosis (Christodoulopoulos, 2025; Nagaraja and Titgemeyer, 2007) and laminitis (Teixeira Passos et al., 2023), and may contribute to the high prevalence of abnormal oral behaviours, such as tongue rolling (Schneider et al., 2020).

Recent anecdotal reports from farmers highlight restlessness, described as intensive, continuous vocalisation, frequent, sustained mounting, and tireless chasing or wild manipulation of feed, as a significant concern affecting bull welfare due to injuries and economic performance due to poor growth and, in some cases, animal losses (Agricultural Chamber Lower Austria, 2024; Federal Information Centre Agriculture, 2024). Suggested causes underlying restlessness include a low proportion of roughage in the feed and mixing of animals (Federal Information Centre Agriculture, 2024). Moreover, in humans, restlessness is a symptom of chronic boredom (Danckert et al., 2018; Finkielsztein, 2024), a highly aversive state often caused by situations characterised by barrenness, which means a lack of stimuli, and monotony, which means a lack of variation – situations many farmed species, including bulls raised for fattening, are faced with.

In bulls, restlessness has only been described anecdotally, but not yet scientifically. However, this behavioural phenomenon has been investigated in other non-human animal species and categories in various contexts. In farrowing sows, for example, restlessness may result from a lack of appropriate substrate and space to show nest-building behaviour (as discussed in Baxter et al., 2024). In dairy cattle, restlessness has been defined as the number of steps (Krebs et al., 2011) or, additionally, weight shifting between legs to study discomfort during forced standing on different surfaces (Chapinal et al., 2011), and as stepping and foot-lifting to compare cows’ behaviour in different automatic milking systems (Gygax et al., 2008). Restlessness has also been defined by an equation that includes lying time, activity and the number of lying bouts in cows around and after parturition (Swartz et al., 2023).

Taken together, some studies have investigated restlessness in pigs and dairy cattle, but it has been defined and recorded in various ways depending on the specific topic of interest. So far, a universal definition of restlessness that allows recording this behavioural phenomenon not just in specific contexts, but more generally, is missing. Moreover, although different studies point to an association between restlessness and some form of negative welfare, research on the potential use of restlessness as an indicator of impaired welfare remains limited.

The general aim of this paper is to gain first insight into restlessness as a potential novel welfare indicator in bulls fattened for meat production. Specifically, we aim to focus on restlessness to answer the following research questions:

1. How can restlessness be operationalised so that it can be quantified?
2. How much does the level of restlessness vary within and across farms, and does it differ between bulls from different weight categories (300, 400, 500 kg)?
3. Does the analysis of behavioural sequences allow for grouping bulls with similar behaviour patterns?
4. How do bulls from different husbandry systems (fully-slatted floor systems, straw-bedded systems, pasture-based systems) differ regarding their level of restlessness?

Research question 1 is conceptual and will therefore be addressed in the methods section, as its definition forms the basis for answering research questions 2, 3, and 4. The latter will be addressed using empirical data from two on-farm studies that build on one another. While our study does not allow us to draw conclusions about the relationship between restlessness and animal welfare concerns, such as (subacute) ruminal acidosis or chronic boredom, we aim to set the stage for future research in this area.

## 2 Animals, Materials and Methods

### 2.1 Animals and husbandry systems

#### 2.1.1 Study 1

This study was conducted on eight commercial bull fattening farms in Upper and Lower Austria that participated in a government-subsidised consulting programme aiming at optimising the profitability of bull fattening farms. The average weight gain of the bulls was 1300 g/day. At least two-thirds of the feed was corn silage. Bulls on all farms were kept on fully-slatted floors; two farms used rubber-covered concrete slats, while the other six farms kept the bulls on concrete slats. On two farms, iron bars were installed horizontally above the bulls to prevent them from mounting. The bulls predominantly originated from dairy farms and were mostly of the dual-purpose breed Fleckvieh.

On average, there were 108 bulls per farm (range: 54 – 160). Group size ranged from four to ten animals per pen, with an average of 7.5 bulls per pen. Space allowance per bull was on average 3.2 m².

#### 2.1.2 Study 2

This study was conducted on eleven commercial bull fattening farms in southern Germany (Bavaria and Baden-Württemberg, n = 9), Thuringia (n = 1) and Schleswig-Holstein (n = 1). The eleven farms differed in their housing systems: 1) fully-slatted floor (FS: n = 4), 2) straw-bedding (SB: n = 4 farms) and 3) organic pasture (OP: n = 3 farms).

The farms were found using personal recommendations and internet research. Based on the criteria breed, age at slaughter, total number of pens with bulls older than 12 months, age difference within a pen, number of bulls per pen, pen size and space per animal, the four most comparable farms per housing system were selected for FS and SB. Since intact bulls kept for fattening purposes are rarely kept on pasture, all thee farms that were identified were included in the study.

##### 2.1.2.1 Fully-slatted floor system

The predominant breed was Fleckvieh or Fleckvieh crosses. Bulls were either raised on the fattening farm or purchased from cattle traders. On average, there were 7.2 bulls per pen (range: 4 – 14), and the average space allowance was 3.5 m^2^ per bull (Table 1). On two farms, iron bars were installed horizontally above the bulls to prevent them from mounting. Fresh feed, provided as a total or partial mixed ration, was provided on average 2.3 times a day. Feed was provided as a total or partial mixed ration. The main feed components were corn silage and grass silage, but the proportions varied between farms. Bulls had enough space at the feeder to be able to eat at the same time. Pens were mostly equipped with a single drinker, with different types used across farms. On average, bulls were 4 months old when housed in this system and slaughtered at an age of 21.7 months.

**Table 1.** Characteristics of the eleven farms included in Study 2.

|  | Fully-slatted | Straw-based | Organic pasture |
| --- | --- | --- | --- |
| Characteristic | Mean | Mean | Mean |
|  | (Min.; Max.; n) | (Min.; Max.; n) | (Min.; Max.; n) |
| Animals per group [n] | 7.2<br>(4; 14; 24<sup>1</sup>) | 20.1<br>(13; 25; 24<sup>1</sup>) | 12.5<br>(4; 18; 6<sup>1</sup>) |
| Space per animal [m<sup>2</sup>] | 3.48<br>(2.45; 5.44; 24<sup>1</sup>) | 5.08<br>(3.62; 8.00; 24<sup>1</sup>) | 2,185<br>(375; 5.000; 6<sup>1</sup>) |
| Ratio of feeding places to animals per pen [n] | 1.1<br>(0.83; 2; 24<sup>1</sup>) | 0.6<br>(0.36; 1; 24<sup>1</sup>) | 1.8<br>(1.38; 2.44; 3<sup>1</sup>) |
| Age during observations per group [months] | 19<br>(16; 22; 21<sup>1</sup>) | 15.6<br>(12; 20.5; 24<sup>1</sup>) | 24.2<br>(19; 33; 6<sup>1</sup>) |
| Age at slaughter [months] | 21.7<br>(20; 23.5; 3<sup>2</sup>) | 18.8<br>(17.5; 22; 4<sup>2</sup>) | 26.3<br>(24; 30; 3<sup>2</sup>) |

<sup>1</sup>: number of pens, <sup>2</sup>: number of farms

##### 2.1.2.2 Straw-bedding system

The predominant breed was Fleckvieh or Fleckvieh crosses. On average, there were 20.1 bulls per pen (range: 13 – 25), and the average space allowance was 5.1 m^2^ per bull (Table 1). Pens consisted of a concrete floor feeding area and a straw-bedded lying area. Two farms provided extra equipment, such as brushes, in the pens. Feed delivery was automated in most farms. On average, feed was provided 3.3 times per day. The main feed components were corn silage and, on two farms, additionally grass silage. The average ratio of feeding places to animals was 0.6. Most pens were equipped with two bowl drinkers. On average, bulls were 3.9 months old when housed in this system and were slaughtered at 18.8 months of age.

##### 2.1.2.3 Organic pasture

All three farms were organic dairy farms that fattened their male offspring on pasture during the summer. With Fleckvieh, Original Swiss Brown and German Black Pied Lowland Cattle, breeds differed between farms. On average, there were 12.5 bulls per pasture (range: 4 – 18), and the average space allowance varied between 375 and 5.000 m^2^ per bull (Table 1). Pasture was managed differently between farms. Farm 1 implemented rotational grazing combined with continuous grazing, Farm 2 provided a sheltered feeding and lying area with permanent access to pasture, and bulls on Farm 3 could continuously graze on one native pasture. The main feed component was grass, either grazed on pasture (Farms 1 and 3) or freshly cut in the sheltered area three times a day (Farm 2). Due to the hot and dry summer and poor growth of grass, haylage was provided in a hayrack during the second farm visit on Farm 3. Pastures were equipped with one trough or bowl drinker. On average, bulls had access to pasture from 1 month until on average 26.3 months of age during the vegetation period.

### 2.2 Study design and data collection

#### 2.2.1 Study 1

Data collection took place in July and August 2020. Each farm was visited twice, with at least 7 days between visits. On each observation day, observations were divided into two sessions: a morning session from 7 a.m. to approximately 1 p.m., and an afternoon session from 2 p.m. to approximately 5 p.m. If the bulls were fed just before the start of the observation, the observation start was postponed for half an hour. The start time of the afternoon recordings was chosen because the animals became more active again after being less active around noon. All observations were conducted by the same observer.

To investigate whether age influences restlessness, bulls across different weight classes were observed, with liveweight serving as a proxy for age. The three different weight classes consisted of animals with approximate liveweights of 400 kg, 500 kg and 600 kg. Bulls were not weighed individually, but their liveweight was estimated by the farmers. On farm 6, only bulls of an average weight of 500 kg were kept.

On each of the eight farms, two pens per weight class, i.e., six pens in total, were selected; on farm 6, six pens of the same weight class were selected. Selection of pens was as much as possible balanced for location within the barn, e.g., pens in the corner versus pens neighbouring two other pens and pens on the left versus pens on the right side of the aisle.

We aimed to observe three bulls from each of the six pens per farm and per observation day, leading to a total of 6 bulls per pen, 36 per farm and 288 in total (36 x 8 farms). However, there was one pen with only five animals and another with only four, reducing the number of observations to 285. Moreover, on some occasions, it was not possible to select focal animals that met our selection criteria, which required animals to be standing or walking (i.e. not lying), but not eating or ruminating, for at least two minutes before the start of the observation. Furthermore, animals looking at the observer before the start of the observation were excluded. Thus, the final number of observations was 270 bulls. Bulls were continuously observed for 15 minutes each, using a tablet equipped with the recording software Mangold Interact®. A folding ladder was used as an observation platform, allowing the observer to be positioned about 1 meter above the ground, thereby ensuring a good view of the bulls, including those in the back of the pens.

The observations of the six focal animals per pen were balanced across morning/afternoon observations, i.e., three bulls were observed in the morning of day 1, and the other three bulls were observed in the afternoon of day 2. To ensure each bull was observed only once, ear tag numbers were recorded. If a focal animal lay down for more than two minutes during the observation, the observation was aborted, and another focal animal was selected unless the animal lay down after 10 minutes, in which case the observation was used.

#### 2.2.2 Study 2

Data collection took place between May and September 2022. Each farm was visited twice, with an average of 65 days between visits, and each visit consisted of two observation days. Observation days were divided into blocks of 2 h, starting with the first block on the first day at 6 a.m. and ending with the last observation block on that day at 8 p.m. Each observation block was followed by a 2-h break. On the second day, the order of observation blocks was reversed, i.e., observations started at 8 a.m. and finished at 10 p.m. As a result, bulls were observed from 6 a.m. to 10 p.m. across the two observation days, totalling 16 hours of observation per farm visit. For the second farm visit, the order of observation and break blocks was reversed. The bulls were observed by the same observer standing on the feed bunk in FS and SB, and outside the pasture equipped with binoculars when needed in OP. Bulls were continuously observed for 8 minutes each, using a tablet equipped with the recording software Mangold Interact®. Observation times were shorter than in Study 1 because continuous observations (data presented here) were complemented by scan samples that covered the activities of all animals in each group (data not shown), which took some time after each individual observation. In total, 722 observations were conducted in FS, 714 observations in SB and 421 observations in OP.

During each farm visit, the three pens with the oldest animals were selected to avoid observation of groups that had recently been mixed. The group with the oldest bulls was selected for OP farms. Within pens, standing or walking bulls were selected according to their position in the pen: FS pens were mentally divided into two equal-sized areas, and the selection of bulls was balanced across the two areas. In SB pens, the feeding and lying areas were mentally divided into two equal-sized areas, and the selection of bulls was balanced across these four areas. Compared to Study 1, bulls in Study 2 were observed repeatedly since we had more observation periods and not enough animals to switch for each observation. The average observation frequency was 4.2 times for FS, 1.5 times for SB and 5.8 times per bull for OP.

### 2.3. Ethogram

#### 2.3.1 Study 1 & 2

We developed an ethogram aiming to capture the full behavioural repertoire of the bulls. The first draft of the ethogram was based on existing ethograms in the scientific literature. It was then complemented by watching existing video clips of bull behaviour and our own direct observations on a farm. The ethogram was then tested and refined during two farm visits in June 2020. The final version of the ethogram comprised 44 behaviours, with some behaviours only being possible in straw-based systems or on pasture (Table 2).

**Table 2.**
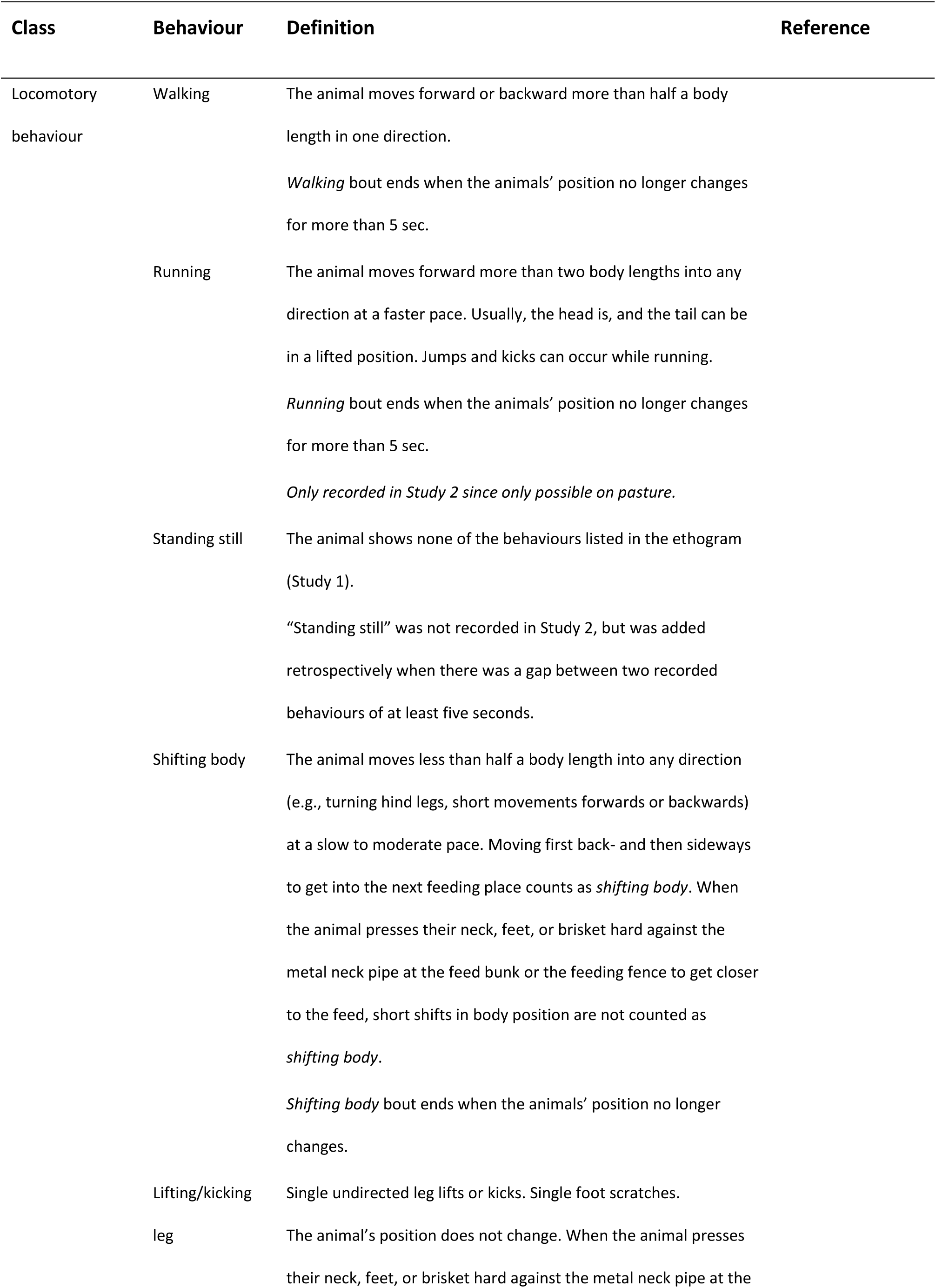
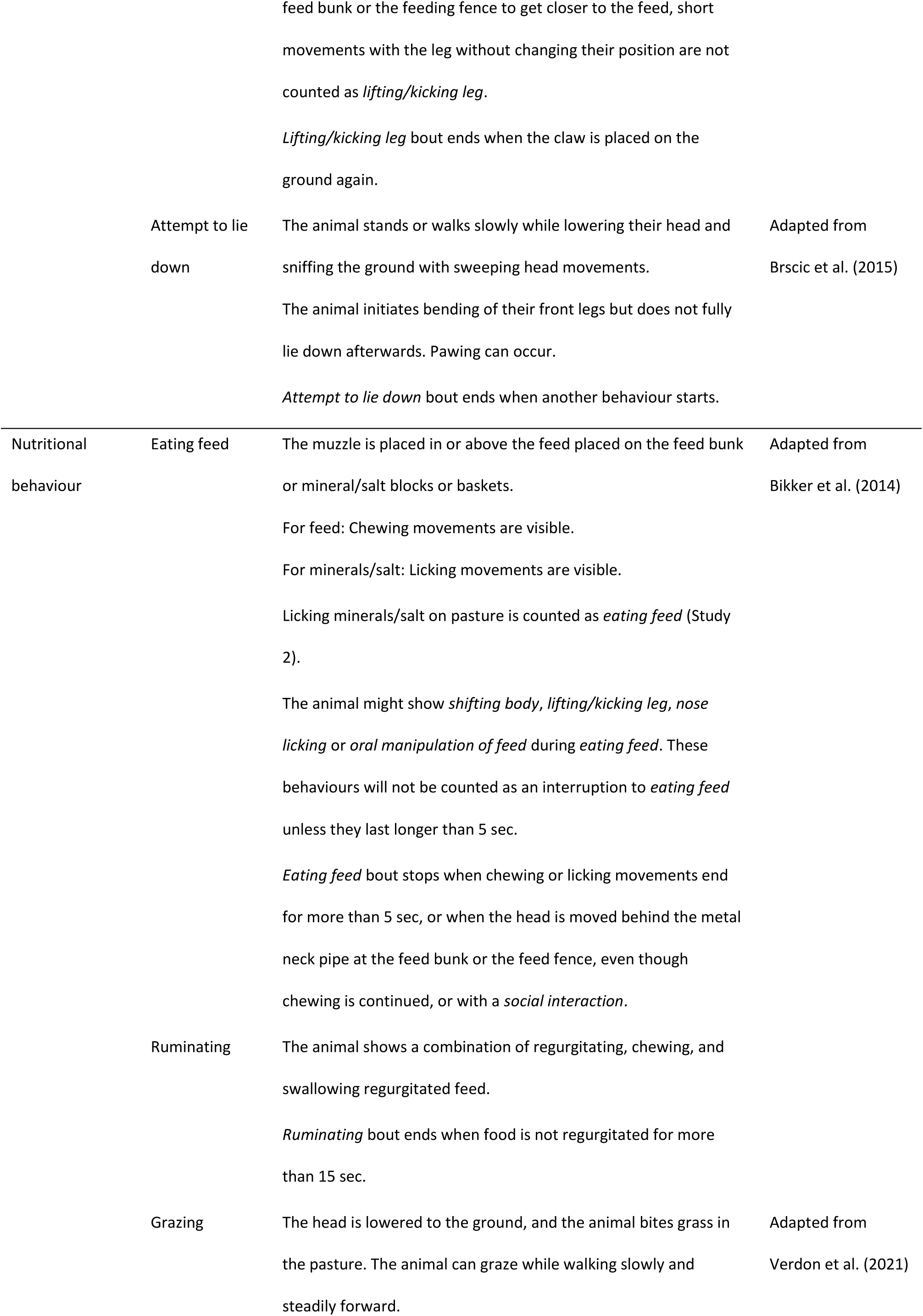
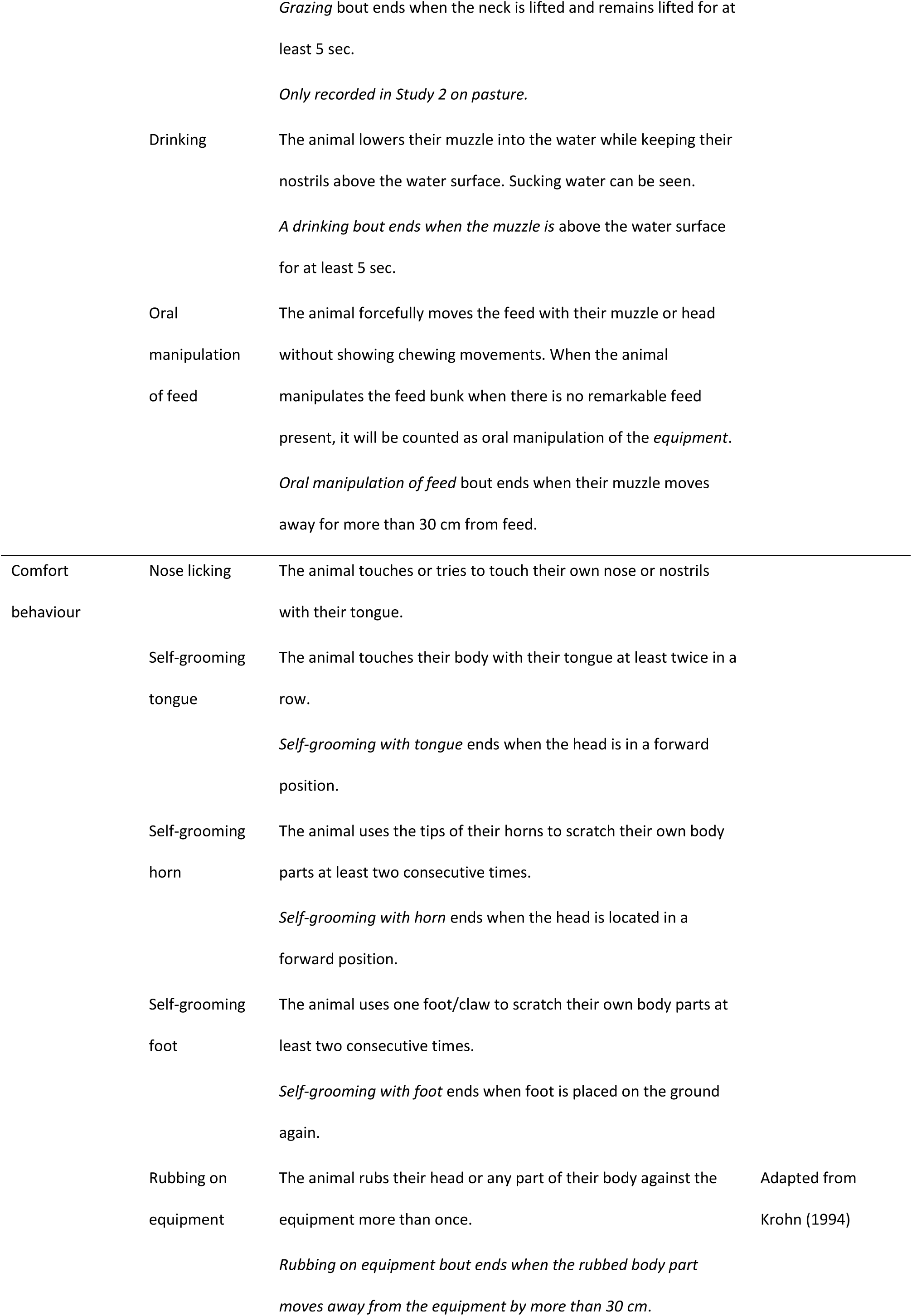
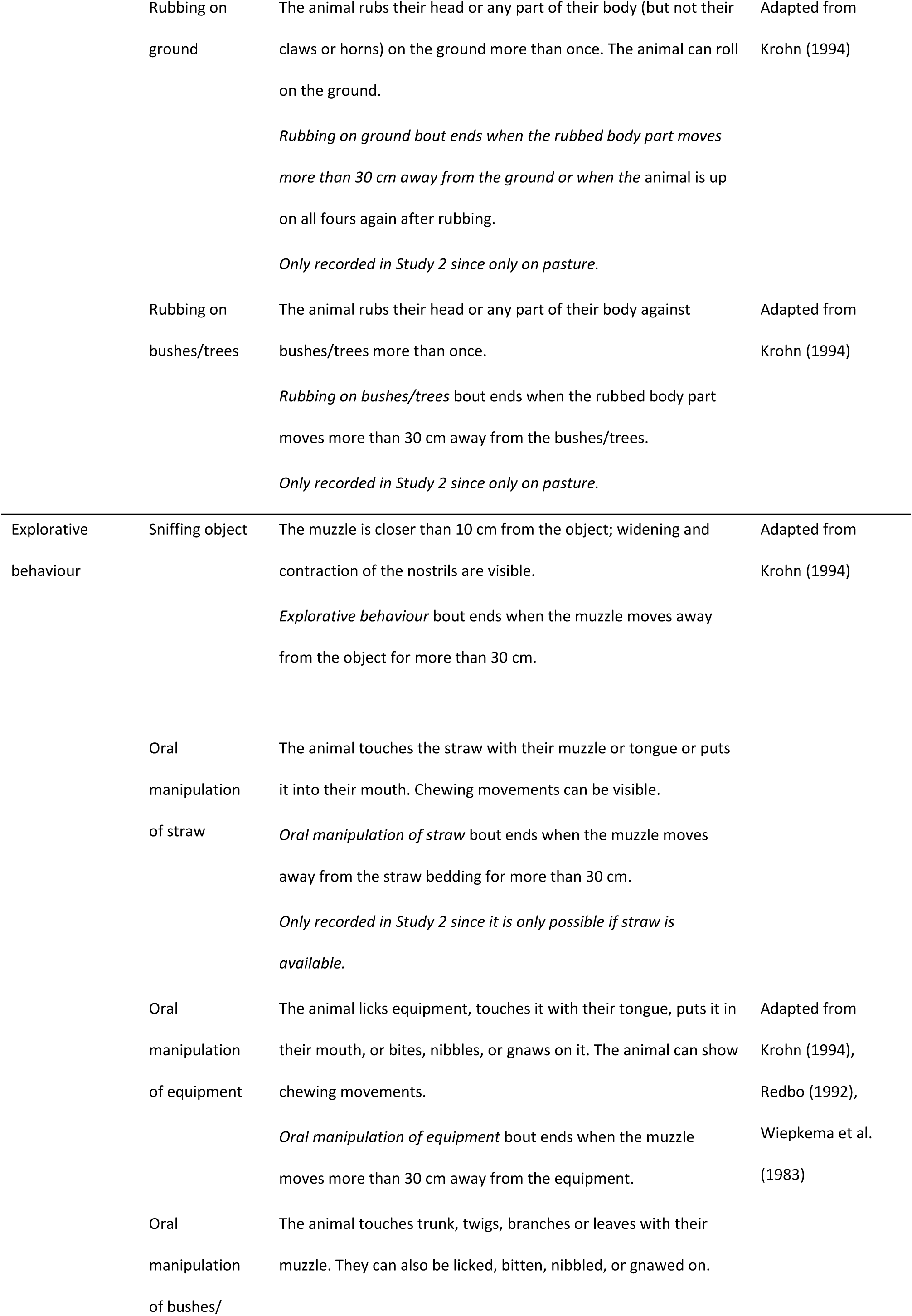
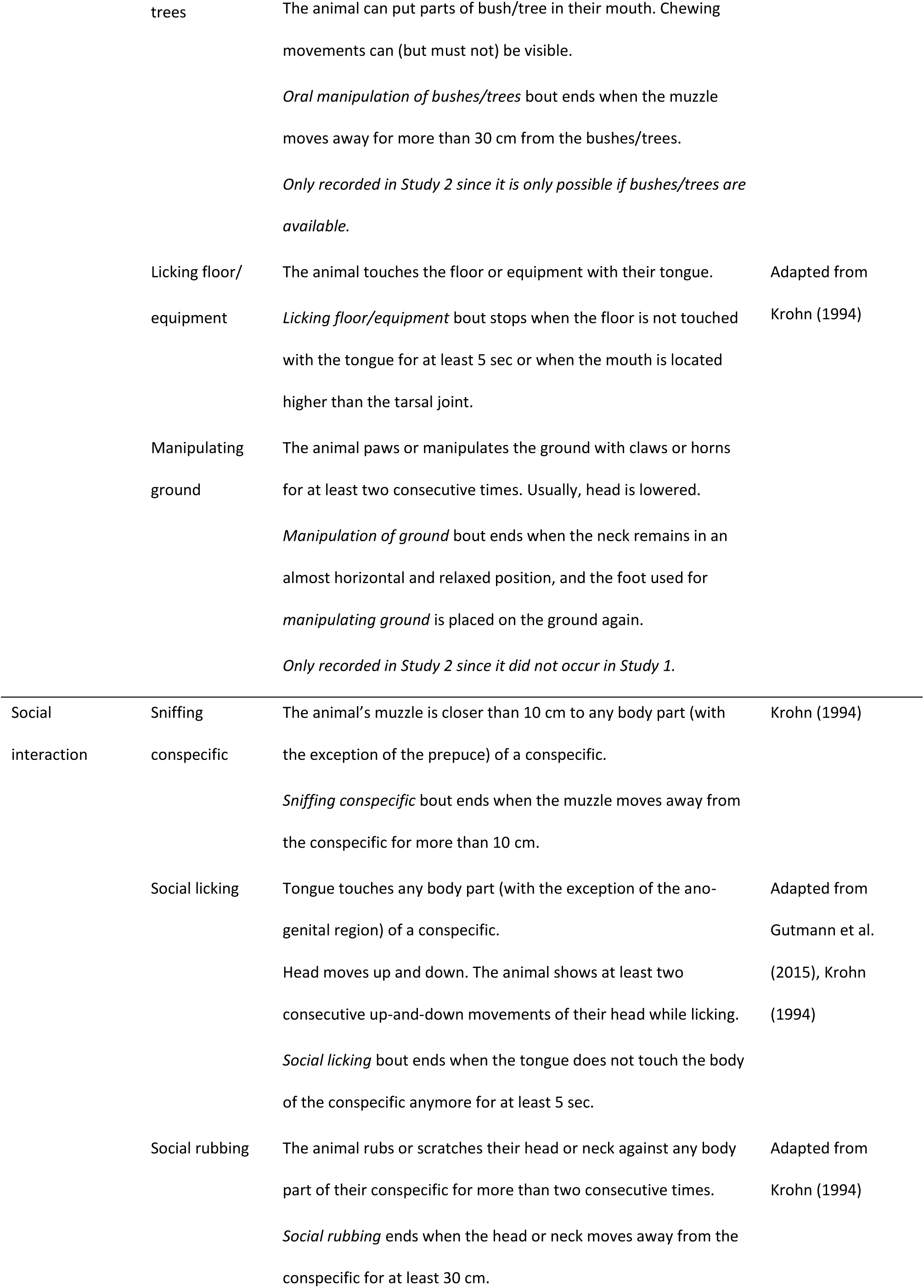
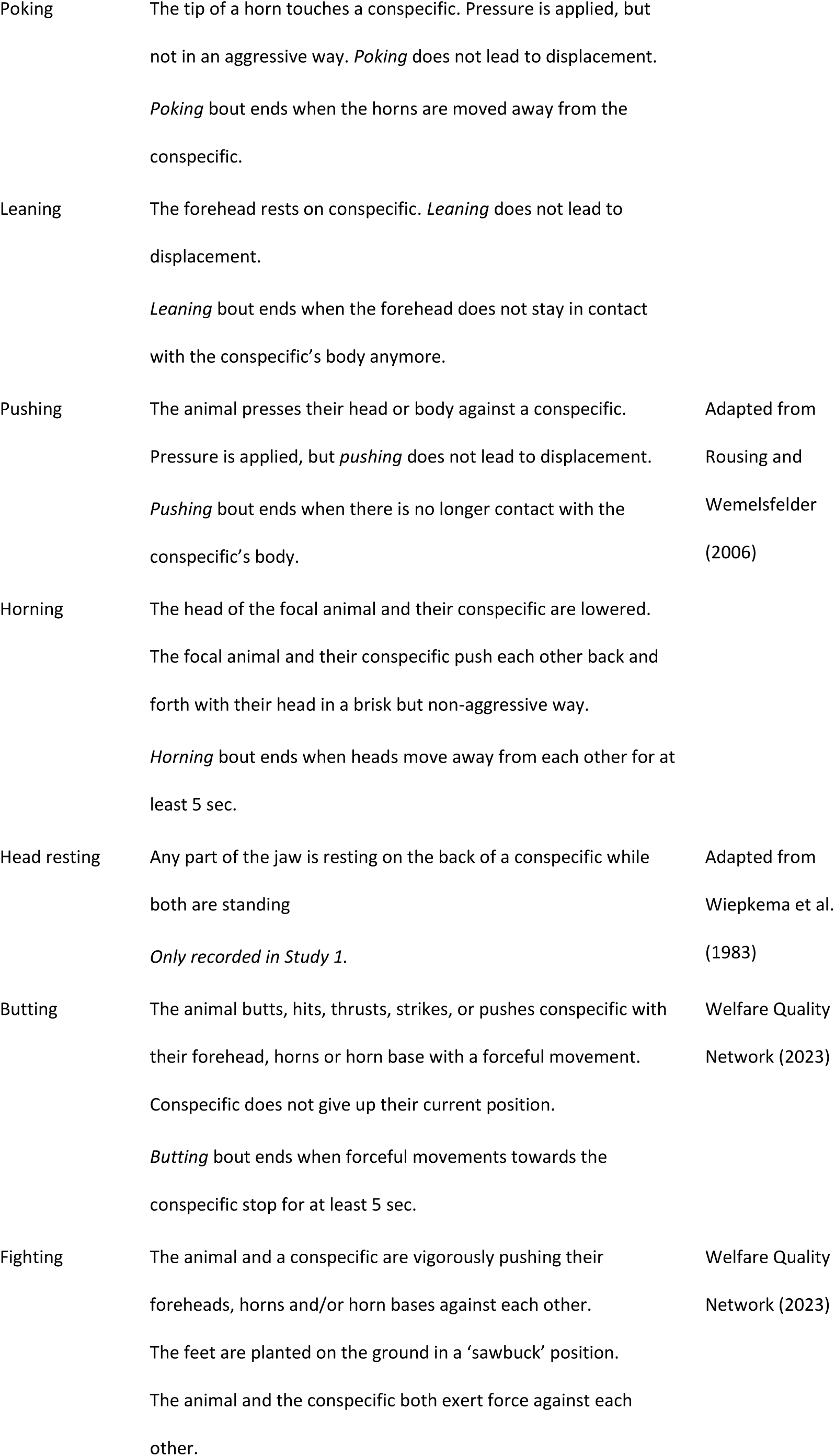
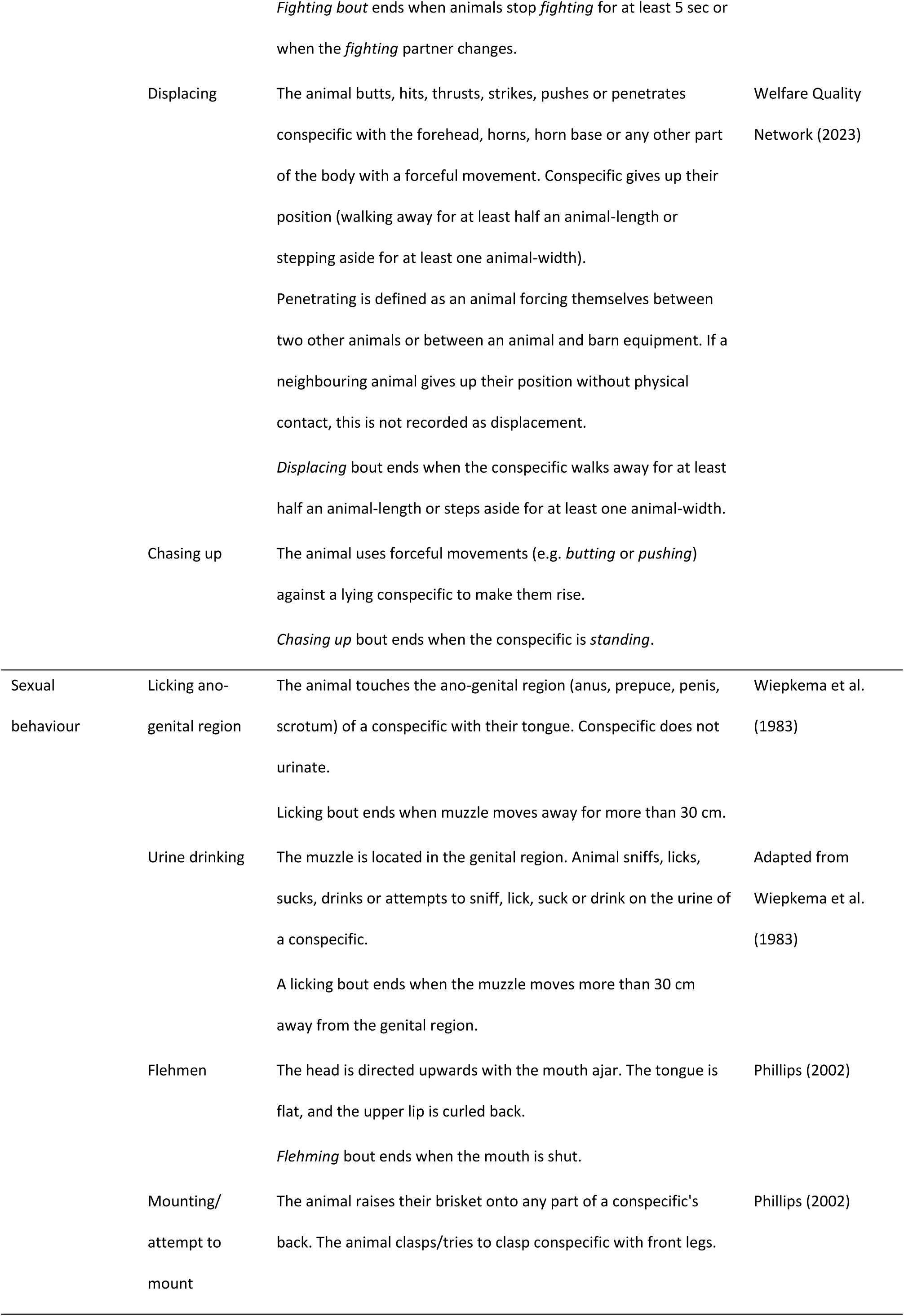
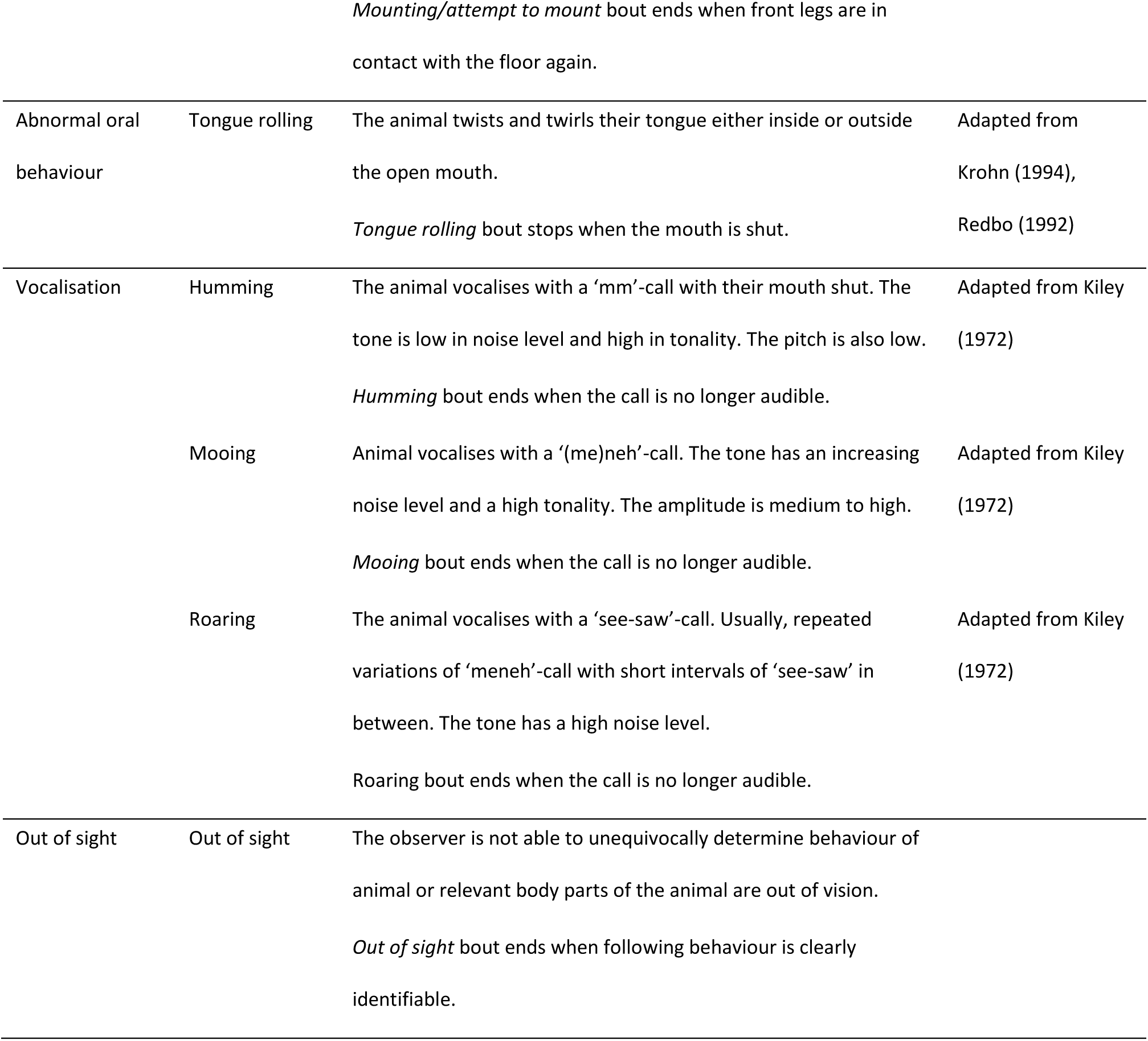
Ethogram of the recorded behaviours in Study 1 and Study 2.

### 2.4 Operational definition of restlessness

Research question 1 addresses the operationalisation of restlessness. Pilot observations revealed that recording durations or frequencies of certain behaviours did not sufficiently capture what we experienced as restlessness. Restlessness is less about the specific behaviours shown and more about the changes between behaviours. We thus operationally defined restlessness by the number of transitions between behaviours in a given time period (Figure 1). The definition is purely quantitative, not distinguishing between transitions between qualitatively different behaviours.

**Figure 1.**
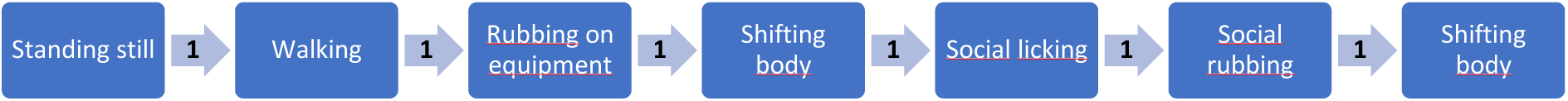
Example of behaviours recorded according to the ethogram (blue boxes) and transitions between these behaviours (arrows between boxes) within a given time frame. This example shows six transitions between behaviours.

### 2.5 Inter-observer agreement

#### 2.5.1 Study 1

To test for inter-observer agreement, focal animals had to meet the same criteria mentioned above, but observation time was reduced to five minutes. On one farm, the behaviour of ten animals kept in five different pens was simultaneously recorded by the observer and another trained observer using Mangold Interact® on two tablets.

#### 2.5.2 Study 2

For inter-observer agreement testing in Study 2, eight randomly selected video recordings from the eight different farms included in Study 1 were used. Per recording, one animal meeting the above-mentioned criteria was selected and their behaviour recorded for ten minutes by two observers independently using Mangold Interact®.

### 2.6 Statistical analyses

All statistical analyses were carried out in RStudio (R 4.1.1. and R 4.5.2).

#### 2.6.1 Study 1

##### 2.6.1.1 Inter-observer agreement

Inter-observer agreement regarding the number of behaviour transitions was assessed by the Intraclass Correlation Coefficient (ICC, function: icc, package: irr) and its 95 % confidence interval. We used a two-way model with ‘agreement’ as ‘type’ and ‘single rater’ as ‘unit’.

##### 2.6.1.2 Data preparation

Since animals sometimes lay down during observations, 61 observations were shorter than 15 minutes. The number of transitions was downscaled or upscaled to a 10-minute observation length. Before calculating the number of transitions, occurrences in which the focal animal was recorded as “out of sight” were removed from the raw data to avoid counting transitions between behaviours and “out of sight” events as transitions. If the animal exhibited different behaviour upon re-entering the field of vision than before the “out of sight” event, a behavioural transition was recorded. If, however, the animal showed the same behaviour after re-entering the field of vision, no behavioural transition was recorded. The total number of “out of sight” events was 577, averaging 2.14 events per observation.

##### 2.6.1.3 Effect of farm and weight class on the number of transitions

We originally planned to run a generalised linear mixed-effects model (function: bglmer, package: blme) with a Poisson family, as the number of transitions is count data. However, due to overdispersion in the outcome measure (variance/mean ratio = 4.32), a Negative Binomial model (family = nbinom1; as using nbinom2 the models did not converge) was fitted instead of a Poisson model. A Likelihood Ratio Test confirmed that the Negative Binomial model provided a significantly better fit than the Poisson model (p < 0.001).

The number of transitions between behaviours served as outcome measure. Fixed effects were farm (categorical, 7 levels; Farm 6 was excluded since there was only one weight class on this farm) and weight class (categorical, three levels, scaled for the models) and their interaction. Random effects were pen nested in weight class nested in farm visit. Farm was not included in the random effect statement because the highest-level effect (i.e., farm) should not be both a random and a fixed effect. To obtain p-values, we ran an ANOVA (base function: anova) to compare the full model, including both fixed effects and their interaction, with a reduced model omitting the interaction or a single main effect. For both fixed effects and the interaction, we used dummy variables with sum contrasts, an approach that allows interpretation of the p-values of the main effects, even when the interaction is significant (Schad et al., 2020).

##### 2.6.1.4 Sequence analysis

In addition to analysing the number of transitions, we aimed to examine the sequence of behavioural transitions to increase our understanding of restlessness. To this end, an unsupervised learning algorithm was used to cluster the sequences of all bulls, taking into account not only the number of transitions, but also the sequences of behaviour, their timing and duration (package: TraMineRextras; (Gabadinho et al., 2011). The sequence analysis was based on the calculation of the “Optimal Matching Distances”. This approach involves measuring pairwise dissimilarities between sequences. The resulting values were used in an unsupervised clustering algorithm (Agglomerative Nesting algorithm = AGNES; (Kaufman and Rousseeuw, 1990) with a dendrogram as the output. Since we were interested in the sequences, down- or upscaling of observation lengths was not possible, so we had to find a compromise between the minimal session length and the number of sessions to include. 249 observations with a length of 580 seconds were included.

To further explore the different clusters, we display the percentage of time bulls spent performing each observed behaviour within each cluster in a stacked bar graph. This illustration does not provide insight into the sequence of behaviours, but it gives an overview of how clusters differ in their time budgets.

#### 2.6.2 Study 2

##### 2.6.2.1 Inter-observer agreement

The Intraclass Correlation Coefficient was calculated as in Study 1.

##### 2.6.2.2 Data preparation

“Out of sight” events were dealt with as described above for Study 1. Since, contrary to Study 1, the behaviour “standing still” was not recorded separately but was relevant for counting transitions as in Study 1, we inserted “standing still” when there was a gap of at least five seconds between two recorded behaviours. This time was chosen to rule out that the time between two consecutive behaviours was due to recording delays. The number of transitions between behaviours was calculated as in Study 1, but downscaled and upscaled to 8 min (not 10 minutes) because of the shorter recording times in Study 2.

##### 2.6.2.3 Effect of husbandry system on the number of transitions

As for Study 1, we originally planned to run a generalised linear mixed-effects model with Poisson as family. However, we also found overdispersion in the outcome measure (variance/mean ratio = 11.37), and thus fitted a Negative Binomial model (family = nbinom2) instead of a Poisson model. A Likelihood Ratio Test confirmed that the Negative Binomial model provided a significantly better fit than the Poisson model (p < 0.001).

The number of transitions served as outcome measure. The only fixed effect was the husbandry system (categorical, three levels). Random effects were pen nested in farm visit day (1,2) nested in farm visit (1,2) nested in farm. Even though we have repeated measures from single individuals, we could not account for this in the random effect statement, as it was not possible to identify individuals, especially in the big groups. The husbandry system was not included in the random effect statement because the highest-level effect (i.e., husbandry system) cannot be both a random and a fixed effect. To obtain p-values, we ran an ANOVA (base function: anova) to compare the full model with the null model (= no fixed effect).

## 3 Results

### 3.1 Inter-observer agreement

Out of 39 possible behaviours, 23 (Study 1) and 21 (Study 2) were observed. The behaviours covered all behavioural classes except for the class “vocalisation”. We covered a range of 26 – 66 transitions in Study 1 and 15 – 56 transitions in Study 2 (both scaled to 10 minutes for better comparison), thereby reflecting both low and high numbers of transitions. Following the framework of Koo and Li (2016), inter-observer agreement for the number of behaviour transitions was ‘very good’ in Study 1 (ICC = 0.84, 95 % CI: 0.46 – 0.96, p < 0.001) and ‘excellent’ in Study 2 (ICC = 0.98, 95 % CI: 0.89 – 0.99, p < 0.001).

### 3.2. Effect of farm and weight class (Study 1)

The average number of transitions between behaviours across farms was 48.3 ± 14.5, with substantial variation within farms (Figure 2). There was an interaction effect between farm and weight class, as well as an effect of farm as main effect on the number of transitions (Interaction: X^2^ = 27.34, p = 0.007, farm: X^2^ = 26.36, p < 0.001; Figure 3). While the number of transitions decreased slightly with increasing weight on Farm 1, it increased slightly on Farm 5. On Farms 2 and 8, the number of transitions was lower in animals weighing approximately 500 kg compared to those weighing 400 kg, and it increased again in heavier animals (600 kg), whereas the pattern was the opposite on Farms 3, 4, and 7. Please note that the pattern description is based on the model estimates, not the medians of the raw data displayed in the boxplots (Figure 3). The main effect of weight class was not statistically supported (X^2^ = 2.34, p = 0.31).

**Figure 2.**
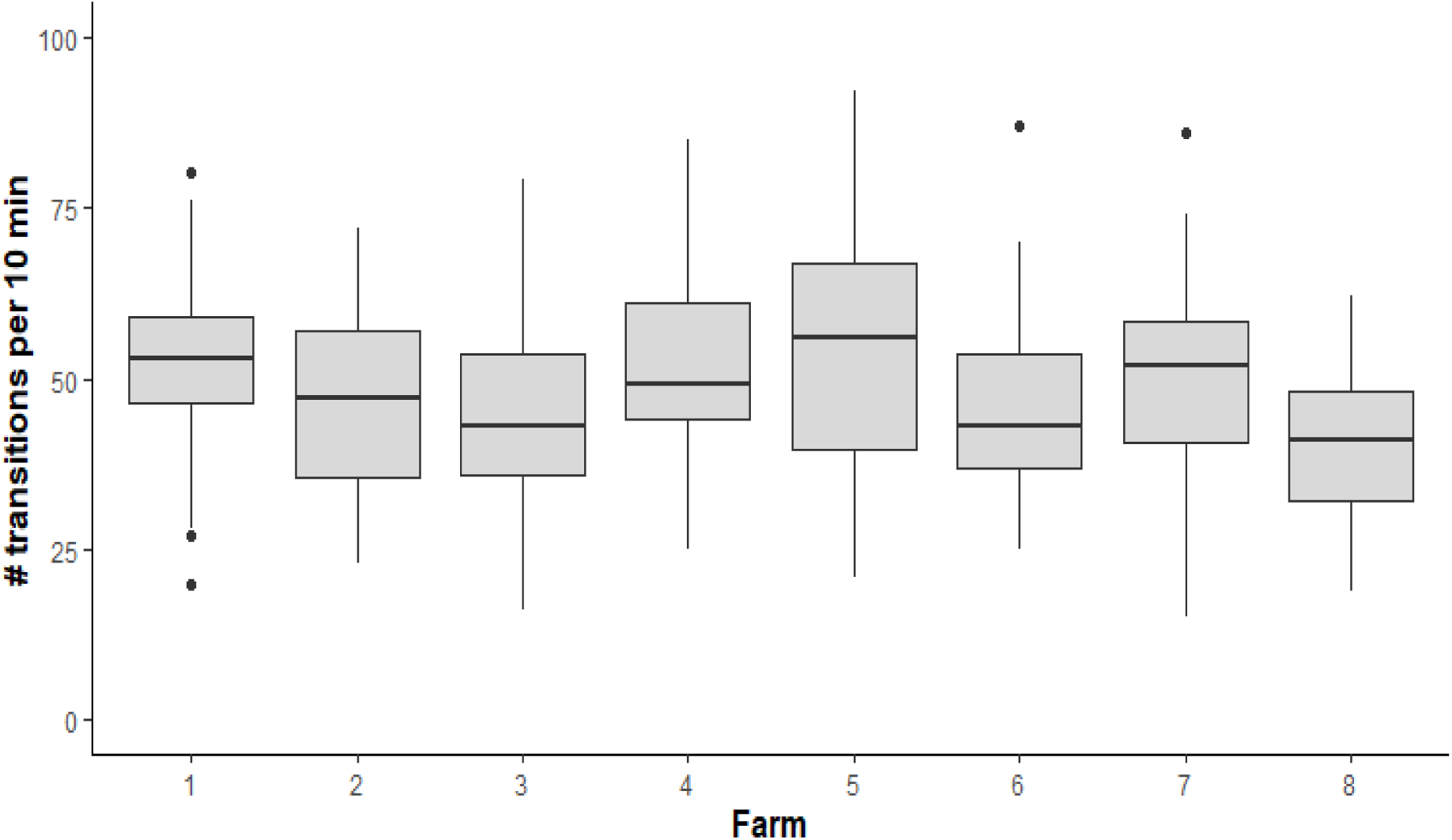
Overview of the individual number of transitions between behaviours per 10-minute observation and the overall variation in transitions between and within the eight farms from Study 1. Boxplots with medians (black line within the box), lower and upper interquartile range (box) and whiskers representing 1.5 times the interquartile range or minimum/maximum values are shown.

**Figure 3.**
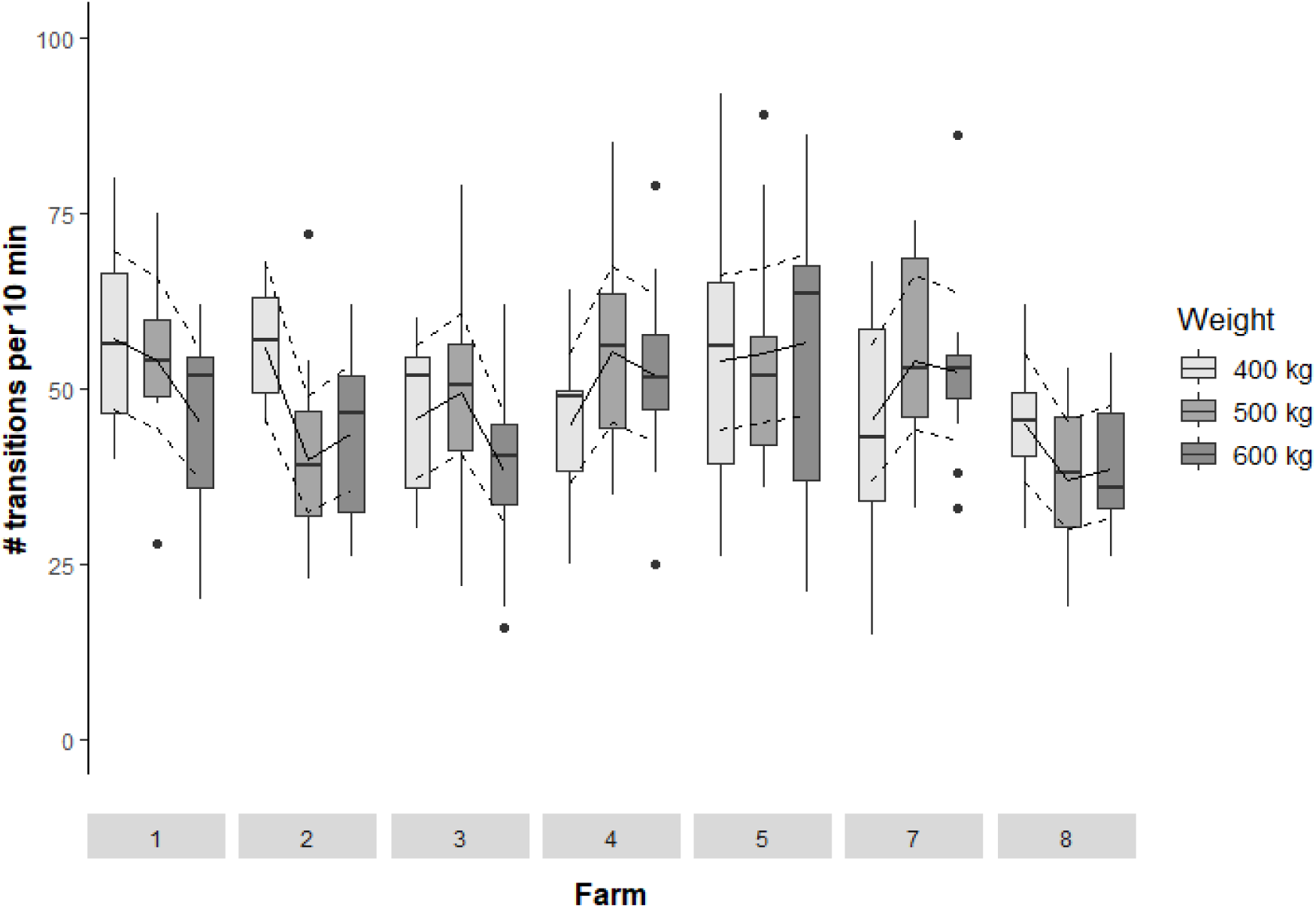
Overview of the individual number of transitions per 10-minute observation split by farm and weight class. Please note that Farm 6 is excluded from this graph since it had only bulls in the weight class of 500 kg, but not 400 kg and 600 kg, at the time of observation. Boxplots with medians (black line within the box), lower and upper interquartile range (box) and whiskers representing 1.5 times the interquartile range or minimum/maximum values are shown. Model estimates and 95 % confidence intervals are drawn as thick and thin lines, respectively.

### 3.3 Sequence Analysis (Study 1)

The sequence analysis revealed an allocation of the 249 observations into four clusters (Figure 4): Cluster 1 with 35 observations and an average of 37 transitions per observation, Cluster 2 with 25 observations and an average of 23 transitions, Cluster 3 with 54 observations and an average of 39 transitions and Cluster 4 with 135 observations and an average of 46 transitions. Figure 5 reveals that across clusters, animals spent most of the observation time standing still or eating, and that the percentages spent in these two behaviours differed: animals in Cluster 3 spent most time standing still, and those in Cluster 4 spent most time eating. Differences between clusters also become apparent when looking at the behaviours, which were performed for less time than eating and standing still. Bulls in Cluster 1 and Cluster 3, for example, spent more time performing such behaviours than bulls in Cluster 2 and Cluster 4, and the percentages they spent performing the different behaviours differed between clusters. Such differences were, here only described exemplarily, a comparably high percentage of social licking in Cluster 3 and of licking and manipulating equipment in Cluster 1.

**Figure 4.**
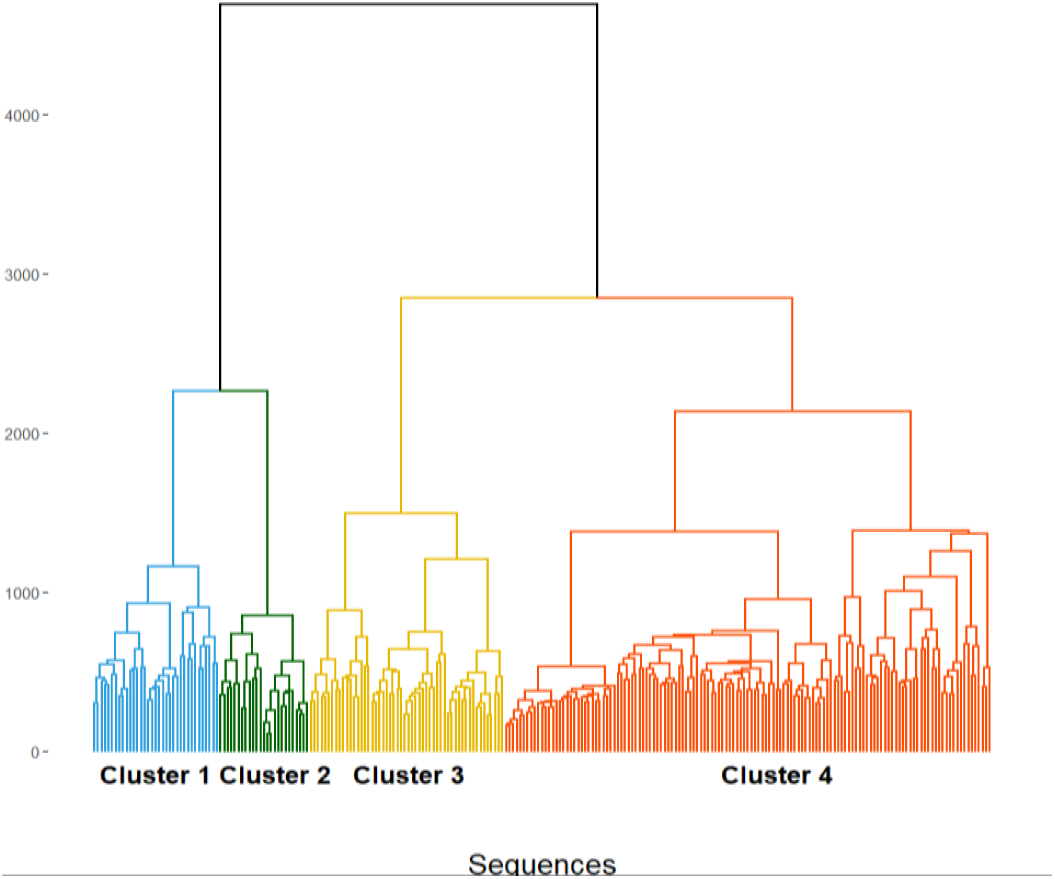
Dendrogram of the clustering process of the sequences of behaviours based on 249 580-second observations. The four different clusters are mirrored by the four different colours. Cluster 2 is the smallest (n = 25 observations), followed by Cluster 1 (n = 35 observations), Cluster 3 (n = 54 observations) and Cluster 4 (n = 135 observations). Cluster 1 and Cluster 2 are most similar to each other, as the link between them is the closest to the bottom of the dendrogram. Clusters 3 and 4 are more similar to each other than they are to Clusters 1 and 2. Please note that irrespective of the absolute values on the y-axis, the relative lengths of the vertical lines indicate the degree of difference, with longer lines representing greater differences.

**Figure 5.**
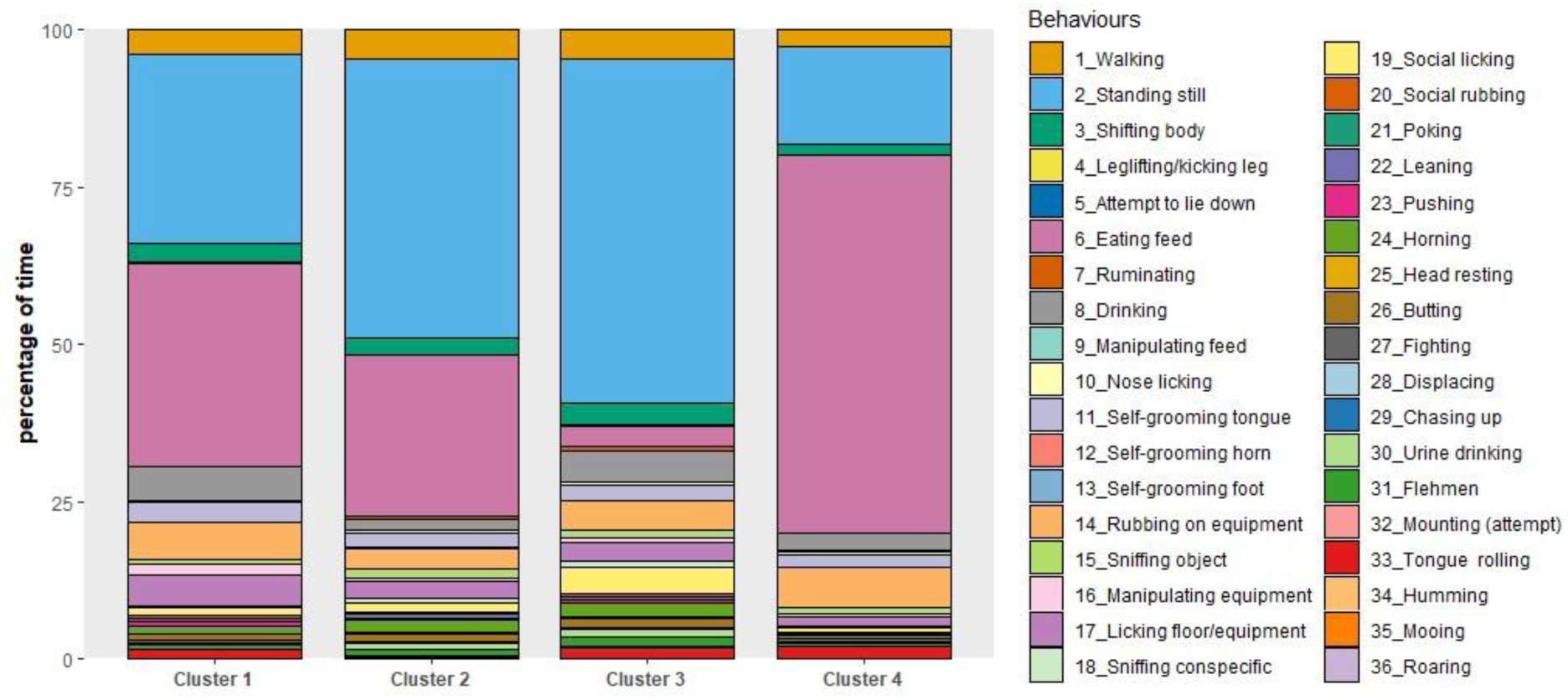
Overview of the percentage of time bulls spent performing each of the recorded behaviours, split by the four clusters.

### 3.4 Effect of husbandry system (Study 2)

The effect of the husbandry system on the number of transitions was statistically supported (X^2^ = 24.2, p < 0.001). The pattern in Figure 6 reveals that the number of transitions was substantially lower in the OP system than in the FS and SB systems, which did not differ from each other. The variation in the number of transitions between farms was smallest in FS systems.

**Figure 6.**
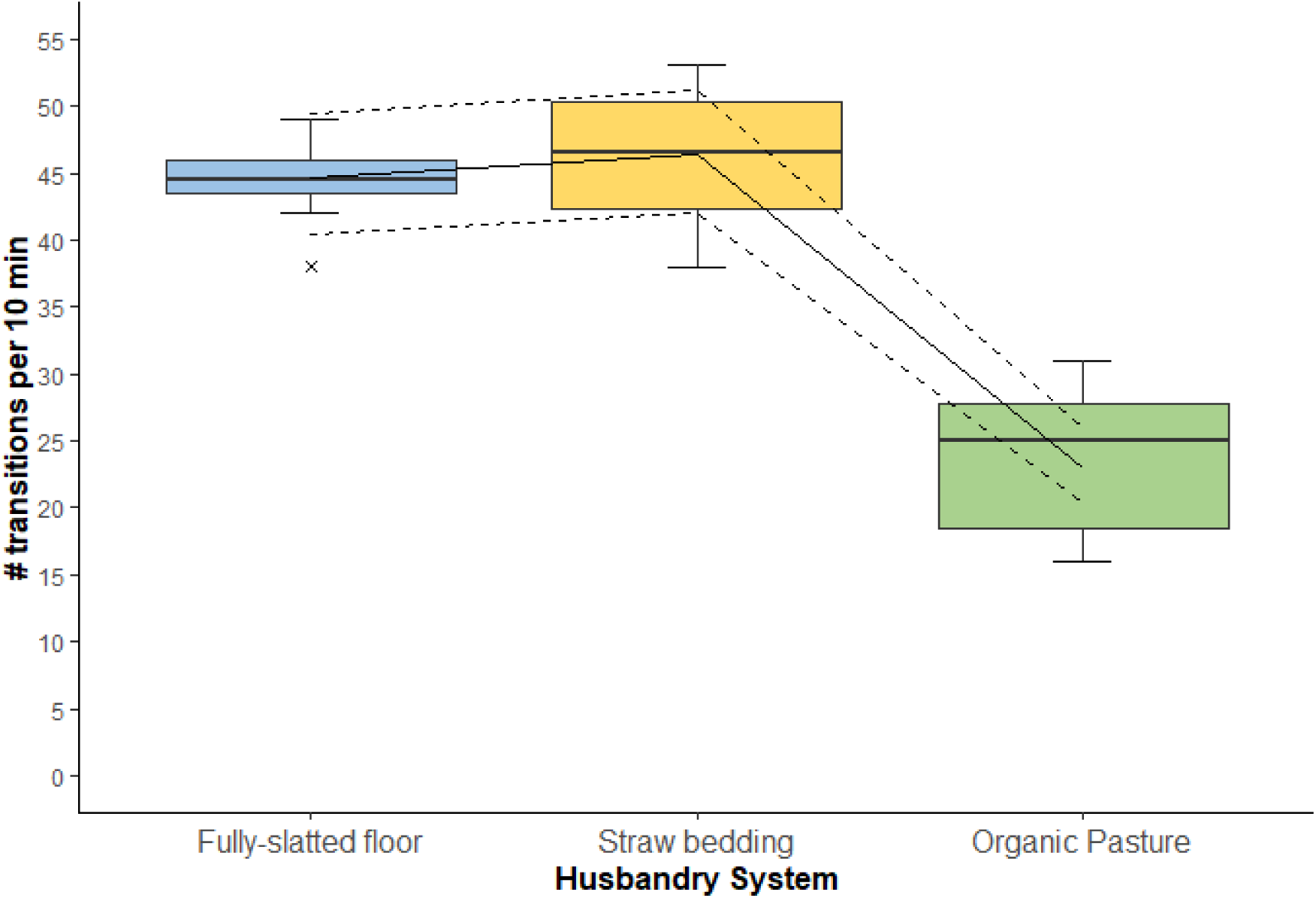
Number of transitions between behaviours per 10-minute observation split by husbandry system, calculated on a farm and farm visit basis. Please note that the number of transitions was upscaled to 10 minutes in this graph to facilitate comparison with Study 1. Boxplots with medians (black line within the box), lower and upper interquartile range (box) and whiskers representing 1.5 times the interquartile range or minimum/maximum values are shown. Model estimates and 95 % confidence intervals are drawn as thick and thin lines, respectively.

## 4 Discussion

This study aimed to provide first insights into the restless behaviour of bulls kept for fattening purposes. We operationally defined restlessness by the number of transitions between behaviours per unit of time. Study 1 revealed an average number of 48.3 transitions per 10 minutes in bulls kept on fully-slatted floors, with a statistically supported difference between farms. Bulls’ weight class (300, 400 or 500 kg) did not affect the number of transitions per se, only in interaction with farm, but there were no clear patterns regarding the direction of effect. The sequence of the recorded behaviours allowed classification of observations into four clusters. Study 2 showed that the husbandry system affects the level of restlessness, with bulls kept on pasture showing fewer transitions than those kept in fully-slatted floor or straw-based systems, whereas the latter two systems did not differ.

### 4.1 Inter-observer agreement

Inter-observer agreement ranged between ‘very good’ on farm (Study 1) and ‘excellent’ on video (Study 2) despite the sometimes very quick transitions between behaviours. On-farm agreement tests reflect the actual recording conditions during data collection, whereas video-based agreement tests have the advantage that videos can be selected to cover a range of conditions. The combined results indicate that the number of transitions can be recorded reliably.

### 4.2 Effect of farm and weight class (Study 1)

With an average of 48.3 transitions within 10 minutes, the results from the eight Austrian farms align well with those from the four fully-slatted floor farms in Germany, where the average was 43.9 transitions within 10 minutes. The number of transitions differed between farms, and we found high variability within and across farms. However, despite this statistically supported effect, the average number of transitions ranged only from 40.4 to 54.7, and there was thus no farm with numbers of transitions as low as, for example, those found on pasture in Study 2, with an average of 23.3 transitions per 10 minutes. While 48 transitions within 10 minutes, which means an average change from one behaviour to another every 12.5 seconds (but with high variation between transition times), seem high, we lack references to classify these numbers as high or low apart from the findings in Study 2. If we consider bulls housed and fed more extensively on pastures as a reference, we can conclude that the number of transitions in the fully-slatted floor system is comparably high. However, we need more research to determine what is “normal” for the number of transitions and, by extension, restlessness, as well as to examine the link between the number of transitions and animal welfare (see also below). Moreover, we excluded animals that were lying or feeding at the start of the observations, so the number of transitions across all animals, including those lying or feeding, may have been lower.

The interaction between farm and weight affected the number of transitions, but we could not identify a clear pattern, so the relevance of this interaction remains unclear. There was no statistical support for an effect of weight class on the number of transitions. This finding may indicate that weight, and related to it bulls’ age, does not affect restlessness levels. However, other explanations also need to be considered. Animals were not weighed using a scale, but weights were estimated by the farmer, an approach prone to error. In addition, as weight was estimated at the pen level, we do not have individual weights to consider in the statistical analysis, which would be more precise than an estimated allocation of groups to classes that differ by 100 kg. Moreover, animals within a pen may differ in weight. Especially when bulls are at the border of two weight groups, the allocation of the whole pen to one weight group may be problematic. Besides this methodological concern, it is possible that while lighter and thus younger animals are more active and thus potentially more restless, the diet of heavier animals is more energy-dense, known to be associated with subacute ruminal acidosis and laminitis (Teixeira Passos et al., 2023), which may contribute to restless behaviour due to pain. Consequently, these potential effects may have offset one another, but this explanation remains speculative. Furthermore, while we found an interaction effect between farm and weight class, no consistent pattern was discernible in farms with obvious differences between weight classes, thus making an interpretation difficult.

### 4.3 Sequence analysis (Study 1)

Based on the sequence of the recorded behaviours, the timing and the average duration of each behaviour, the 249 observations could be allocated to four clusters that differed in size, with respect to the number of observations per cluster, the number of transitions, as well as the time spent performing different behaviours. Our results provide insights into the temporal structure and representation of behaviour types, including, but not limited to, the number of transitions, but we need to acknowledge that animals were not always in comparable states. In Cluster 4, for example, animals spent more time eating than in the other clusters, making a comparison difficult. Yet, across clusters, the relative distribution of observations as a function of time of the day was similar, indicating that changes in behaviour patterns typically observed throughout the day did not affect the type of behavioural sequences. We thus consider our approach promising, but currently still too premature to identify what constitutes truly restless behaviour patterns.

Illustrating the percentage of time the bulls spent performing different behaviours does not necessarily explain the differences between clusters, which are based on the sequence of behaviours, but may still help better characterise the clusters. Bulls spent most of their time eating and standing still, but with highly varying proportions across clusters. One important note when interpreting the results is that standing still does not necessarily mean that the bulls were standing still for a longer period. In contrast, standing still was also recorded as a transition between other behaviours when none of the recorded behaviours was performed. In future studies, we propose to differentiate between standing still for a predefined period, which can be interpreted as inactivity (as, for example, in Hintze et al. 2020 for periods over 30 seconds), and short transitions between different behaviours. Moreover, the relatively short observation periods provide only a snapshot of the bulls’ behaviour, and the single observations do thus not characterise the individuals. As this was the first study focusing on restlessness in bulls, we decided to record transitions across farms and among many animals to gain initial insight into variation between farms and animals. We suggest that future studies should include longer and more observations per individual to classify individual bulls according to their behaviour.

### 4.4 Effect of the husbandry system (Study 2)

Bulls on pasture showed fewer transitions between behaviours than bulls in indoor systems, indicating that they were potentially less restless than their conspecifics on FS and SB farms. In addition to the pronounced differences in housing conditions between pasture farms and both indoor systems, FS and SB, feeding also differed greatly, with bulls on pasture only being fed roughage, but no concentrates. Both differences in housing and feeding, or their combination, may have contributed to the lower number of transitions on pasture. While the difference between the pasture farms and the indoor farms was quite pronounced, we need to consider that we had fewer animals to observe, that the animals were, on average, a bit older, and that the pasture farms kept different breeds. These confounding factors could not be excluded, as they are inherent to the pasture-based system, but they need to be considered when interpreting the results. Moreover, we observed animals repeatedly but could not account for this in the statistical models since it was impossible to identify each individual in the large pens on SB farms and on pasture. Thus, there is a risk of pseudoreplication, which needs to be kept in mind when interpreting the results. Variability was lower on FS farms than on SB and OP farms. This may be explained by the more standardised FS systems and the bigger system differences on SB and OP farms.

Interestingly, the level of restlessness did not differ between bulls on FS and SB farms. SB pens are more structured than FS pens and can thus be considered less barren. However, the level of monotony, i.e., lack of variation, may be similarly high in both systems, providing a potential, though speculative, explanation for why restlessness did not differ between husbandry systems. Another (or additional) explanation may be grounded in the comparably high feeding intensity in both systems. High levels of rapidly digestible carbohydrates in the feeding ration are known to lower the ruminal pH, which may result in (subacute) ruminal acidosis (Christodoulopoulos, 2025). Even though the mechanism remains unclear, rapidly fermentable carbohydrates are also known to cause laminitis, likely due to endotoxins entering the bloodstream in case of acidosis, when the ruminal mucosa is morbid (Teixeira Passos et al., 2023). Both acidosis and laminitis are associated with discomfort and pain, which, in turn, may cause restlessness. While bulls on SB farms had access to straw as bedding, FS bulls received physically effective fibre through grass silage during seven out of the eight farm visits. Grass silage was provided to SB bulls only during three of the eight farm visits. Thus, while housing conditions may be considered more intensive in FS than in SB, feeding intensities were similar, if not more intensive, on SB farms. Consequently, it is possible that the monotony in the housing conditions, the feeding intensity or a combination of both resulted in the higher number of transitions in FS and SB compared to OP farms, but these explanations remain speculative. Future research should disentangle variation in monotony level and feeding intensity using a factorial experimental design to better understand the causes of restlessness.

### 4.5 Association of behavioural transitions and animal welfare (Studies 1 & 2)

Since this is the first study to investigate restlessness across the whole behavioural repertoire of cattle, we do not know what a “good” or “normal” number of transitions is. One could argue that bulls on pasture could be seen as a reference system for “what is normal”, but we lack independent information on the welfare status of these animals, and, in addition, could not avoid inherent confounding factors between the systems, rendering it difficult to draw strong conclusions about the association between restlessness and welfare.

Our operational definition of restlessness is quantitative rather than qualitative, meaning it is based on the number of transitions between behaviours over a given period, independent of the meaning of the different behaviours for the bulls. Consequently, one may criticise that the level of restlessness of animals transitioning between negatively connotated behaviours may be the same as that of animals transitioning between rather positive behaviours. However, we believe that quick transitions between behaviours indicate restlessness, independent of the behaviours’ valence. Quick transitions between positively connotated behaviours, e.g., one lick of a conspecific or one bite of feed, that are then interrupted by other behaviours, may not be perceived as positive. While one advantage of our definition may be that it is independent of the difficult classification of which behaviours are positive and which are negative (and in which context, Keeling et al., (2021), we currently lack knowledge of what a good or natural duration of a behavioural bout would be.

Moreover, restlessness should not be confused with behavioural diversity, since the latter focuses on the diversity of behaviours displayed (Miller et al., 2020), whereas restless animals may show a higher number of transitions between only a few different behaviours. We propose that future studies develop an analytical approach that considers both the number of transitions, the diversity, as well as the qualitative nature of different behaviours.

## 5 Conclusion and outlook for future research

Restlessness is a behavioural phenomenon in bulls kept for fattening purposes that farmers have anecdotally described as a potential welfare and economic problem, but which has so far received no attention in research. With this study, we provide first insights into restlessness by offering an operational definition and describing the level of restlessness across commercial farms and husbandry systems, as well as different weight groups. However, our study does not allow us to draw conclusions about the welfare consequences related to restlessness. Future studies should examine the relationship between restlessness and behavioural diversity, focus on understanding the causes of restlessness, and illuminate what this behavioural phenomenon means for bull welfare.

## Acknowledgments

We would like to thank all bull farmers for their openness in welcoming us to observe their animals, thus supporting science and increasing our knowledge about this category of cattle.

